# Does increased gait variability improve stability when faced with an expected balance perturbation during treadmill walking?

**DOI:** 10.1101/2020.10.19.345934

**Authors:** Jacqueline Nestico, Alison Novak, Stephen D. Perry, Avril Mansfield

## Abstract

**Background:** Currently, there is uncertainty as to whether movement variability is errorful or exploratory.

**Research question:** This study aimed to determine if gait variability represents exploration to improve stability. We hypothesized that 1) spatiotemporal gait features will be more variable prior to an expected perturbation than during unperturbed walking, and 2) increased spatiotemporal gait variability pre-perturbation will correlate with improved stability post-perturbation.

**Methods:** Sixteen healthy young adults completed 15 treadmill walking trials within a motion simulator under two conditions: unperturbed and expecting a perturbation. Participants were instructed not to expect a perturbation for unperturbed trials, and to expect a single transient medio-lateral balance perturbation for perturbed trials. Kinematic data were collected during the trials. Twenty steps were recorded post-perturbation. Unperturbed and pre-perturbation gait variabilities were defined by the short- and long-term variabilities of step length, width, and time, using 100 steps from pre-perturbation and unperturbed trials. Paired t-tests identified between-condition differences in variabilities. Stability was defined as the number of steps to centre of mass restabilization post-perturbation. Multiple regression analyses determined the effect of pre-perturbation variability on stability.

**Results:** Long-term step width variability was significantly higher pre-perturbation compared to unperturbed walking (mean difference=0.28cm, p=0.0073), with no significant differences between conditions for step length or time variabilities. There was no significant relationship between pre-perturbation variability and post-perturbation restabilization.

**Significance:** Increased pre-perturbation step width variability was neither beneficial nor detrimental to stability. However, the increased variability in medio-lateral foot placement suggests that participants adopted an exploratory strategy in anticipation of a perturbation.

## Introduction

There are two opposing interpretations of variability during motor tasks. The first describes variability as a central nervous system error that interferes with achieving motor goals and assumes that errors should be minimized to achieve optimal performance [1]. This interpretation is supported by studies showing increased movement variability in people with neurodegenerative conditions, such as Parkinson’s or Huntington’s diseases, compared to people without these conditions, which interferes with optimal motor task performance [2]. Furthermore, older adults exhibit increased gait variability, which is often used as an indicator of increased fall risk [3].

The opposing interpretation argues that movement variability is an exploratory behaviour, which allows for rich sensory input from the environment [4]. To ensure the reliability of neural networks, it is best to optimize errors and noise, which are unavoidable, and learn to retrieve motor memories in their constant presence [5]. Thus, rather than attempting to supress errors and noise, the brain learns to exploit them to reliably store memories [5].

The exploratory nature of variability has been observed with saccadic eye movements, which provide variable visual input, allowing individuals to gather rich sensory information from their surrounding environment [6]. This exploratory role of movement variability has also been demonstrated for balance tasks. When centre of mass (COM) motion was reduced via a ‘locking’ mechanism, significant centre of pressure (COP) movement variability was still observed [4]. When participants expected an external postural perturbation to quiet standing they increased COP movements [7], which may be a strategy to increase sensory input and improve the response to the perturbation. Indeed, increased COP and COM variability in quiet standing prior to a perturbation improved stability, as indicated by the need to take a step after the perturbation [8]. Combined, these studies suggest that movement variability during quiet standing allows for sensory exploration as a strategy to maintain stability [4].

The aforementioned studies [4,6,8] provide support for an exploratory role of movement variability during quiet standing. To our knowledge, no study to date has examined the potential exploratory role of movement variability during human gait. The potential for gait variability to enhance stability is evident in studies examining legged robots, which were originally designed with a set number of walking patterns [9]. With minimized movement variability, the robots would become unstable and be unable to adjust their balance in challenging environments or in response to perturbations [9]. Newer technology has allowed robots to be programmed with variable walking patterns that draw upon multiple degrees of freedom [10] so they can better adapt to such situations [9].

The purpose of this study was to determine if movement variability is a strategy to explore the environment and improve stability when walking. We define variability as changes in spatio-temporal gait parameters (step length (SL), step width (SW), and step time (ST)) from one step to the next and stability as the ability to respond to a perturbation [8]. We hypothesized that: 1) spatio-temporal gait parameters will be more variable prior to trials when a balance perturbation is expected than when no perturbation expected; and 2) increased spatio-temporal gait variability prior to the perturbation will correlate with improved stability after the perturbation.

## Methods

### Participants

Sixteen young, healthy adults (20-35 years old, eight women), volunteered to participate in this study. Participants were excluded if they had neurological or sensory impairment, or prior balance training. Written informed consent was obtained from all participants. The study protocol was approved by the University Health Network Research Ethics Board (ID# 18-6326).

### Apparatus

Data were collected within the Challenging Environment Assessment Laboratory at the Toronto Rehabilitation Institute, which contains a motion platform, capable of moving in all directions (Figure 1). For this study, a lab space measuring 5.6 × 5.1 × 3.8m was configured onto the motion platform. A treadmill was fixed into the lab space, with the handrails removed. Participants wore a safety harness that was secured to the ceiling of the lab overhead.

**Figure 1:**
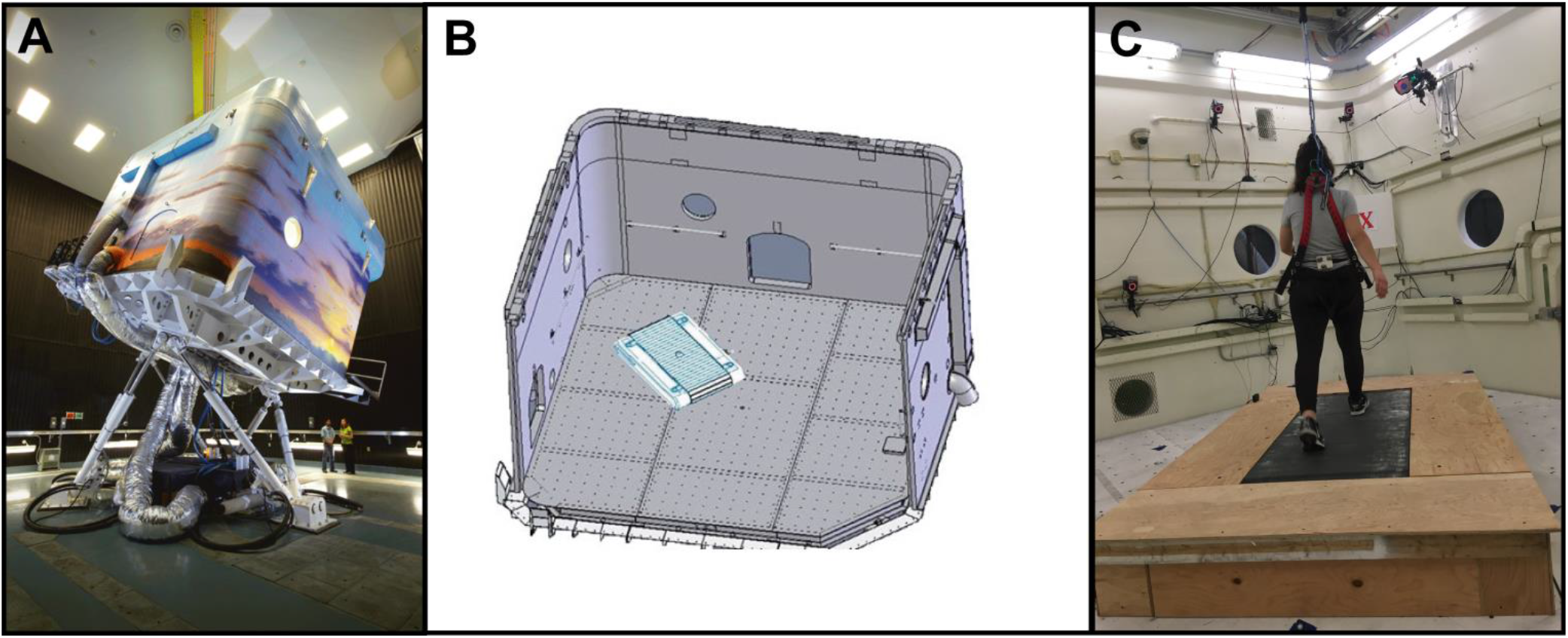
Experimental set-up. A) Outside of the motion base with the laboratory pod fixed on it. B) Schematic of the inside of the lab space with the treadmill fixed to the floor. C) A participant on the treadmill, wearing a harness. A red ‘X’ was placed in front of the treadmill to provide a focal point for participants to focus on during all trials.

Participants were outfitted with reflective markers secured to their left and right heels, base of first metatarsals, lateral malleoli, fifth metatarsal phalangeal joints, and tip of the first phalanges, and a cluster placed on the lower back. The lab space was equipped with 15 motion capture cameras (11 x Kestrel 2200 cameras & 4 x Raptor-E, recording using Cortex 6.2.9.1745, Motion Analysis Corp., Santa Rosa, California, USA), which collected data at a frequency of 200Hz.

The motion platform delivered medio-lateral (ML) translational perturbations to destabilize participants and evoke balance reactions during treadmill walking. The perturbations followed a waveform of 200ms acceleration followed immediately by 200ms deceleration. The peak displacement, velocity, and acceleration were 0.072m, 0.36m/s, and 1.8 m/s^2^, respectively.

### Protocol

Data collection occurred within a single session. Participants’ height and weight were measured, and limb dominance was assessed. The International Physical Activity Questionnaire (IPAQ [11]) and Endler Multidimensional Anxiety Scale – Trait (EMAS-T; [12]) were administered at the start of the session. Treadmill walking speed was set according to the average of three 10-m overground walking trials. Participants were then outfitted with the motion capture equipment and completed a brief familiarization. The familiarization consisted of 1 minute of walking to acclimate to walking on the treadmill within the lab space, and four perturbations (2 from the left and 2 from the right) while standing with the treadmill stationary so that participants would know what to expect of the perturbations.

Participants completed 15 treadmill-walking trials (Table 1), each lasting up to two minutes, presented in two blocks: 1) unperturbed walking (no perturbation expected); and 2) expected perturbation. In the unperturbed walking block, participants were told that the motion platform would not move. The expected perturbation block consisted of 11 trials; participants were told that the platform would likely move during these trials. Participants were instructed to recover their balance as naturally as possible and continue walking if a perturbation occurred. During eight of the perturbed trials, perturbations occurred 75-105 seconds after the start of the trial (i.e., ‘late perturbation’); this allowed us to collect at least 100 steps of steady-state walking prior to each perturbation to reliably estimate spatio-temporal variability [13]. The treadmill stopped and the trial ended when the participant had taken at least 20 steps after the perturbation. Two trials had early perturbations (15-45 seconds after the start of the trial) and one did not have a perturbation; these trials were used to ensure participants did not always anticipate the perturbation occurring towards the end of the trial. All perturbations were manually triggered by a research assistant during single support.

**Table 1:**
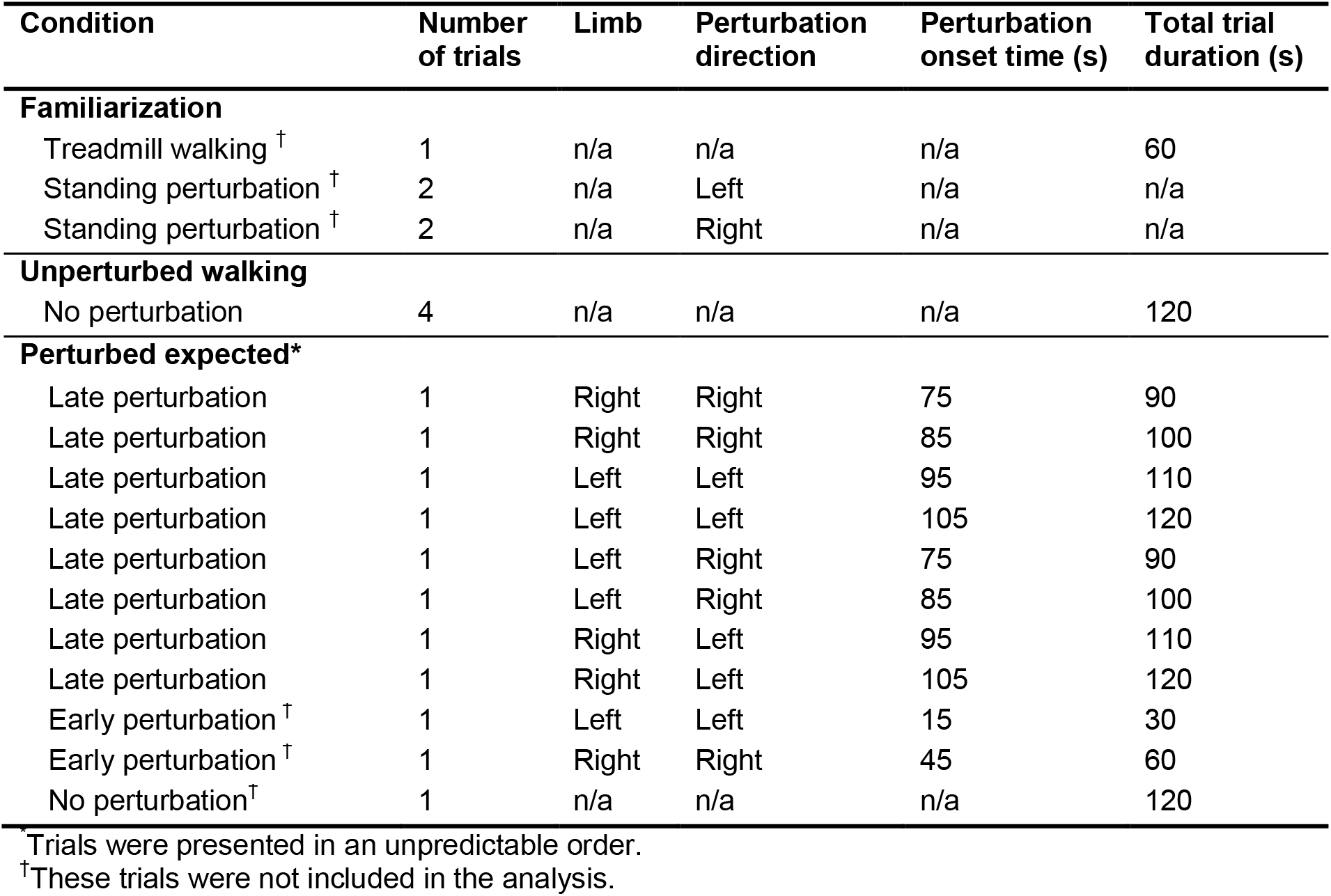
Trial conditions. The limb indicates the stance limb when the perturbation occurred; all perturbations were triggered to occur during single support.

| Condition | Number of trials | Limb | Perturbation direction | Perturbation onset time (s) | Total trial duration (s) |
| --- | --- | --- | --- | --- | --- |
| **Familiarization** |  |  |  |  |  |
| Treadmill walking <sup>†</sup> | 1 | n/a | n/a | n/a | 60 |
| Standing perturbation <sup>†</sup> | 2 | n/a | Left | n/a | n/a |
| Standing perturbation <sup>†</sup> | 2 | n/a | Right | n/a | n/a |
| **Unperturbed walking** |  |  |  |  |  |
| No perturbation | 4 | n/a | n/a | n/a | 120 |
| **Perturbed expected\*** |  |  |  |  |  |
| Late perturbation | 1 | Right | Right | 75 | 90 |
| Late perturbation | 1 | Right | Right | 85 | 100 |
| Late perturbation | 1 | Left | Left | 95 | 110 |
| Late perturbation | 1 | Left | Left | 105 | 120 |
| Late perturbation | 1 | Left | Right | 75 | 90 |
| Late perturbation | 1 | Left | Right | 85 | 100 |
| Late perturbation | 1 | Right | Left | 95 | 110 |
| Late perturbation | 1 | Right | Left | 105 | 120 |
| Early perturbation <sup>†</sup> | 1 | Left | Left | 15 | 30 |
| Early perturbation <sup>†</sup> | 1 | Right | Right | 45 | 60 |
| No perturbation <sup>†</sup> | 1 | n/a | n/a | n/a | 120 |

\*Trials were presented in an unpredictable order.
<sup>†</sup>These trials were not included in the analysis.

The order of trial blocks was counterbalanced between participants. After each block, participants were asked if they believed the researcher when they were told the platform would or would not move. The Endler Multidimensional Anxiety Scale – State (EMAS-S [12]) was also administered following each condition.

### Data processing

Data were initially processed using Visual3D (Version 6.03.6 C-Motion, Germantown, Maryland, USA). Marker positions were filtered using a fourth order dual-pass zero phase lag Butterworth filter with a cut-off frequency of 6 Hz. Left and right heel marker velocities were used to determine timing of heel strikes. Heel strike was defined as the time after toe-off when heel marker velocity was first < 0.1 m/s. SL and SW were defined as the absolute anterior-posterior (AP) and ML distances, respectively, between the heel markers at heel strike. ST was defined as the absolute difference between the time of consecutive heel strikes. Custom MATLAB routines (R2014a, The Mathworks, Inc., Natick, Massachusetts, United States) were used for the remainder of data processing. Perturbation onset time was defined as the time when the motion base acceleration exceeded 0.1m/s^2^.

### Outcomes

#### Short- and long-term variability

Poincaré plots were used to calculate short- and long-term spatio-temporal variability during each trial. For each trial, each point in the time series (i.e., *x(n+1)*) was plotted as a function of the previous one (i.e., *x(n)* [14]). The data points were positioned about a line with a slope of 1 passing through the origin. Two standard descriptors, SD1 and SD2, were calculated as follows [14]:

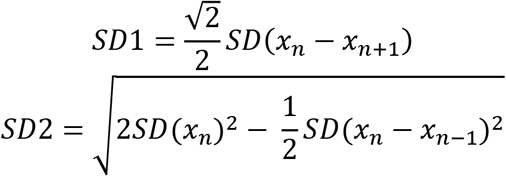

Where SD is the standard deviation of the time series. SD1 represents short-term, or step-to-step, variability, while SD2 indicates long-term variability, which is the continuous variability over the course of the trial when step-to-step variability is removed [15].

#### Number of steps to COM velocity restabilization (stability)

ML COM velocity was calculated at each heel strike. Steps to recovery was defined as the number of steps between the perturbation and when the absolute ML COM velocity entered and remained within a predefined threshold for at least four consecutive steps (Figure 2). The threshold was defined as the mean ±3 standard deviations of the absolute ML COM velocity at heel strike for 100 steps prior to the perturbation.

**Figure 2:**
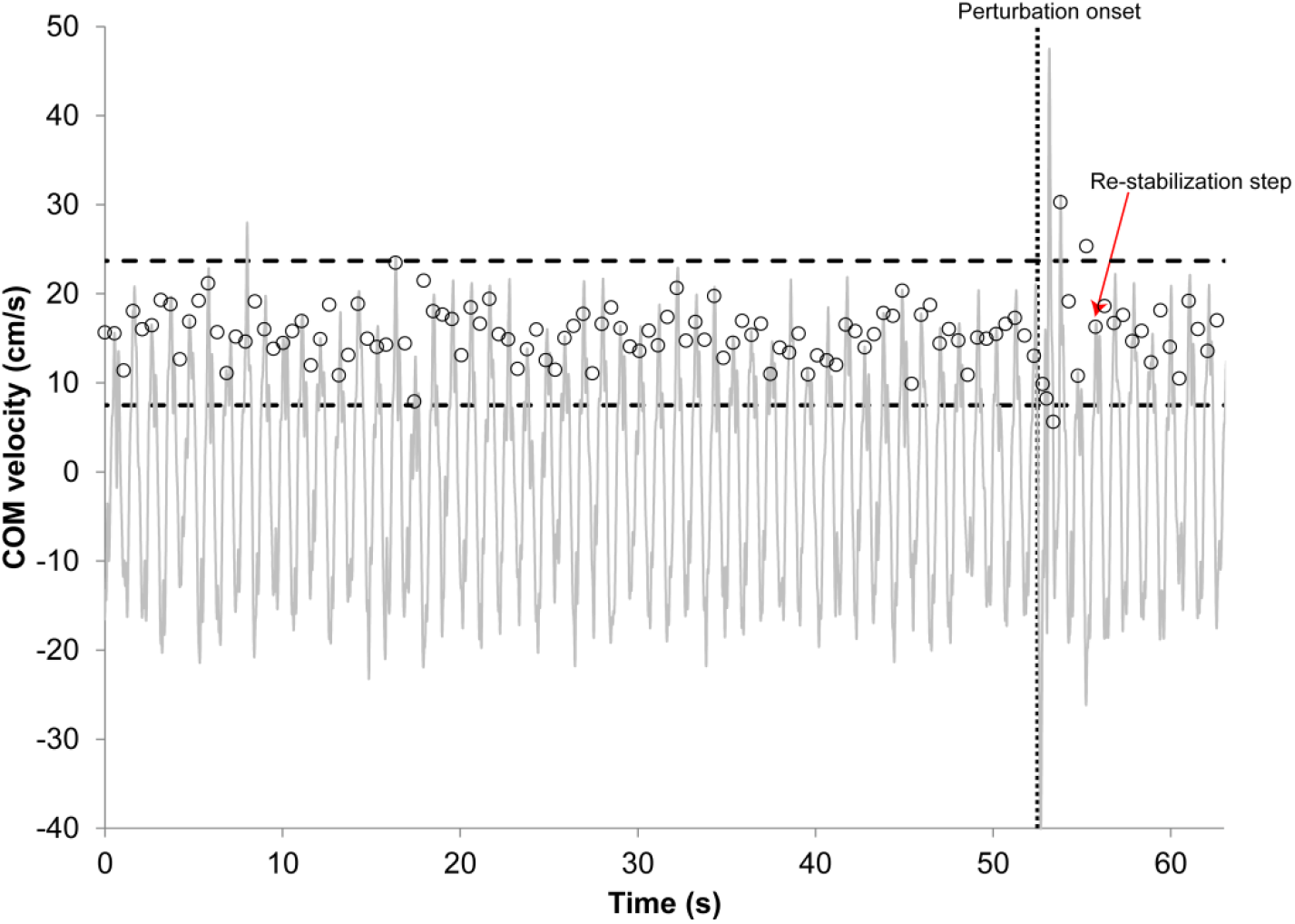
Steps to restabilization. The grey line shows the medio-lateral (ML) centre of mass (COM) velocity throughout the trial. Open circles show the absolute ML COM velocity at heel strike for each step. For simplicity, only the 100 steps prior to the perturbation are shown. The dashed horizontal lines show the upper and lower bound of the absolute ML COM velocity in the 100 steps prior to the perturbation (mean ± three standard deviations). The dotted vertical line is the perturbation onset time. The restabilization step, indicated with a red arrow, is the first step where the absolute ML COM velocity remains within the pre-perturbation threshold for at least four consecutive steps.

### Statistical analysis

Only the unperturbed walking and late perturbation trials were included in the analysis. Perturbed trials had two components: 1) PrePERT, which refers to the 100 steps pre-perturbation, where participants anticipated a perturbation, and 2) PostPERT, which refers to steps following the perturbation. Unperturbed trials had one component: NoPERT, which refers to the 100 steps from the middle of the unperturbed walking trials where participants did not anticipate a perturbation.

To test the first hypothesis, paired t-tests were used to compare short- and long-term spatio-variability (SL, SW, ST) between PrePERT and NoPERT. A Holm-Bonferroni correction [16] was used to adjust for multiple comparisons with the initial alpha value set at 0.0083. To test the second hypothesis, multiple regression with generalized estimating equations and repeated measures was used to determine the relationship between short- and/or long-term spatio-temporal variabilities and steps to regain stability. The time of the perturbation onset after heel strike (expressed as a percentage of a ‘typical’ unperturbed gait cycle) and the mean SW PrePERT were additional potential confounding variables included in the model. Two sets of general linear models were conducted to account for the perturbation direction (i.e., toward the stance limb or swing limb). Testing of the second hypothesis focused only on variabilities that differed between conditions (NoPERT and PrePERT), with alpha being adjusted accordingly. An auto-regression correlation structure was applied to the model, to account for changes over time from trial to trial.

Additional analyses were conducted to help explain the results. Paired t-tests were used to compare the average SL, SW, or ST between PrePERT and NoPERT. Paired t-tests were also used to determine differences in EMAS-S scores between PrePERT and NoPERT. Spearman correlations were used to determine the relationship between EMAS-S scores and PrePERT variability. Alpha was set to 00.05 for these additional analyses.

## Results

All participants reported that they believed that the platform would not move during Block A and that it would likely move during Block B. Long-term SW variability was significantly higher PrePERT compared to NoPERT (t_15_=−3.10, p=0.0073, Figure 3). There were no significant between-condition differences in short-term SL (t_15_=0.03, p=0.98), SW (t_15_=−1.42, p=0.18), ST (t_15_=1.28, p=0.22), or long-term SL (t_15_=0.09, p=0.93) and ST variabilities (t_15_=0.24, p=0.82; Figure 3).

**Figure 3:**
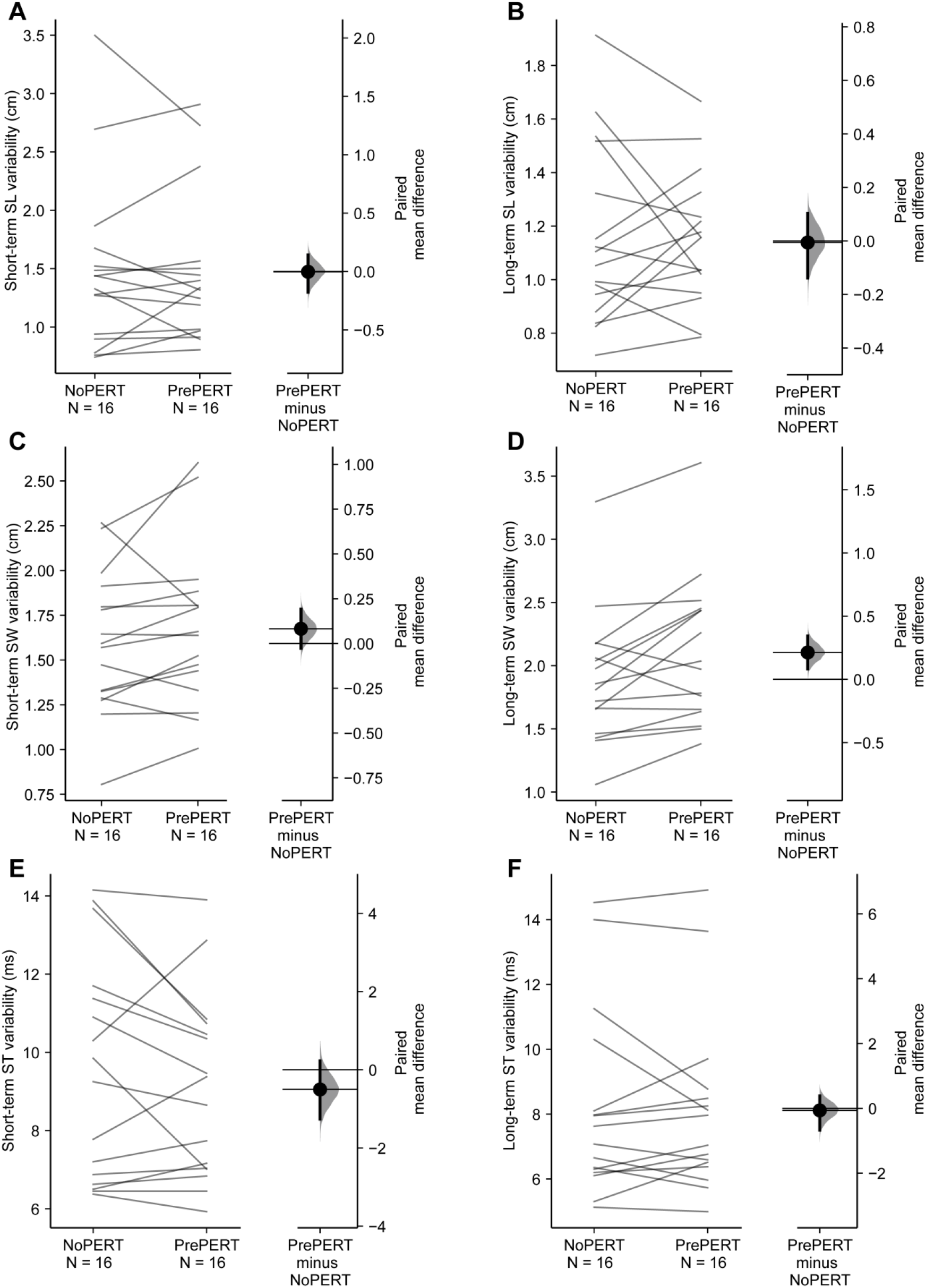
Spatio-temporal variability between conditions. A) Short-term step length (SL) variability, B) long-term SL variability, C) short-term step width (SW) variability, D) long-term SW variability, E) short-term ST variability, and F) long-term step time (ST) variability PrePERT compared to NoPERT. The paired mean differences between NoPERT and PrePERT is shown in the above Gardner-Altman estimation plots. Both groups are plotted on the left axes as a slopegraph: each paired set of observations is connected by a line. The paired mean difference is plotted on a floating axes on the right as a bootstrap sampling distribution. The mean difference is depicted as a dot; the 95% confidence interval is indicated by the ends of the vertical error bar [17]. Only long-term SW variability (D) was significantly greater PrePERT compared to NoPERT.

Perturbed walking trials where perturbations occurred during double support (30/128) were removed from analysis of the second hypothesis. Of the remaining 98 trials, 52 had the perturbation directed towards the stance limb and 46 had the perturbation directed towards the swing limb. Participants took 1-12 steps to restabilize ML COM velocity PostPERT. Since long-term SW variability was the only variability measure that exhibited differences between conditions, the multiple regression analysis focused on the relationship between long-term SW variability and restabilization PostPERT. Alpha was set to 0.025. When accounting for the additional variables, the percent of the gait cycle in which the perturbation occurred and mean SW PrePERT, there was no statistically significant relationship between long-term SW variability PrePERT and the number of steps to stability PostPERT when the perturbation was directed towards the stance (estimate=−1.39, p=0.15) or swing limb (estimate=0.18, p=0.65; Table 2).

**Table 2:** **Results of multiple regression analysis.** of the effect of increased long-term step width variability PrePERT on the number of steps to ML COM velocity restabilization PostPERT.

| Perturbation direction | Variable | Estimate | Standard error | 95% confidence limits |  | Z | p |
| --- | --- | --- | --- | --- | --- | --- | --- |
| Towards swing limb | SD2 SW (cm) | -0.18 | 0.38 | -0.93 | 0.58 | -0.45 | 0.65 |
|  | GaitCycle (%) | 0.006 | 0.17 | -0.27 | 0.38 | 0.36 | 0.72 |
|  | MeanSW (cm) | 9.18 | 14.28 | -18.82 | 37.17 | 0.64 | 0.52 |
| Towards stance limb | SD2 SW (cm) | -1.39 | 0.97 | -3.30 | 0.52 | -1.43 | 0.15 |
|  | GaitCycle (%) | 0.072 | 0.24 | -0.39 | 0.54 | 0.31 | 0.76 |
|  | MeanSW (cm) | -22.44 | 5.73 | -33.67 | -11.21 | -3.92 | <.0001 |

GaitCycle= percent in gait cycle that perturbation onset occurred
SD2= long-term variability

Mean SW was significantly higher PrePERT compared to NoPERT (t_15_=−2.74, p=0.015; Figure 4). There were no significant differences between SL or ST variabilities PrePERT compared to NoPERT (t_15_=0.94, p=0.36; t_15_=1.46, p=0.17, respectively). Endler Multidimensional Anxiety Scale – State (EMAS-S) scores from fifteen participants were included in this analysis; one participant’s scores were not useable due to human error. EMAS-S total scores were significantly higher following the expected perturbation condition (mean score: 25.8, standard deviation: 7.3) compared to the unperturbed condition (mean score: 36.7, standard deviation: 15.8; t_14_>3.60, p<0.0021). There was no significant correlation between the EMAS-S total score and long-term SW variability PrePERT (r=0.26, p=0.36).

**Figure 4:**
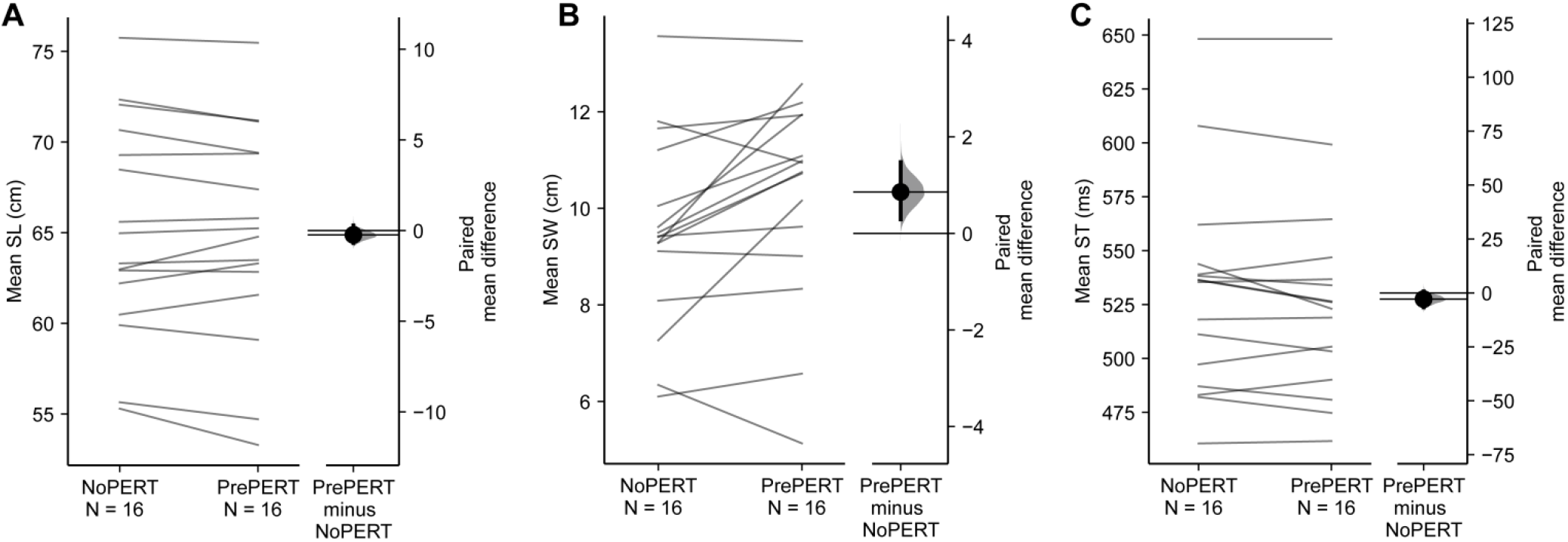
Average spatio-temporal measures between conditions. A) step length (SL), B) step width (SW), and C) step time (ST). The paired mean differences between NoPERT and PrePERT is shown in the above Gardner-Altman estimation plots. Both groups are plotted on the left axes as a slopegraph: each paired set of observations is connected by a line. The paired mean difference is plotted on a floating axes on the right as a bootstrap sampling distribution. The mean difference is depicted as a dot; the 95% confidence interval is indicated by the ends of the vertical error bar [17]. Participants walked with significantly wider steps PrePERT compared to NoPERT.

## Discussion

The purpose of the present study was to better understand if variability during gait has a functional role for stability. We hypothesized that participants would have more variable gait in the steps leading up to an anticipated medio-lateral perturbation compared to during unperturbed walking. Additionally, we hypothesized that pre-perturbation variability would increase stability. We found that 1) participants had higher long-term step width variability when expecting a perturbation compared to when they did not; and 2) increased long-term step width variability prior to the perturbation did not influence stability.

While there was no evidence that increased long-term SW variability prior to the perturbation influenced stability (i.e., the number of steps to restabilization after the perturbation), the increased long-term SW variability may have had a role in participants’ ability to perform the task. From a dynamical systems theory perspective, increased variability can ensure a system does not become too static in a complex environment so that functional movement solutions can be applied during exploratory behaviours [18]. This aligns with research suggesting that there is an optimal state of variability allowing for flexible movements [19], reflecting a healthy system’s ability to adapt to the environment [20], and that being outside of this range is indicative of unhealthy movements [20]. For example, studies have found that individuals with certain neurological conditions (e.g., Parkinson’s and Huntington’s diseases) have increased spatio-temporal gait variability [2]. Likewise, increased spatio-temporal gait variability is related to increased risk of falling in older adults [3] and is considered a marker of poor dynamic balance control [21]. In addition, increased postural movements are known to arise when individuals anticipate a perturbation [7,22,23], and have been suggested to limit the effect of the perturbation to the whole body [7,22]. It has also been suggested that spatio-temporal variability is a feature of young, healthy adults’ ability to adapt during perturbed walking [24]. Here, decreased variability would indicate inflexible motor behaviours with less adaptability in changing environments, while increased variability indicates a noisy, unpredictable system [20]. Taken together, perhaps the increased long-term SW variability observed when anticipating a perturbation in the present study was simply within a healthy, optimal range and enabled participants to be more flexible in selecting an appropriate response. Future studies should determine how movement variability in anticipation of a perturbation changes with age and neurologic disease.

Another possible explanation for our findings is that increased long-term SW variability reflected increased attentional demands of walking due to anxiety when anticipating a perturbation [23]. Indeed, EMAS-S scores revealed that anxiety was higher when expecting a perturbation compared to when no perturbation was expected, and long-term SW variability and EMAS-S were positively correlated, although this correlation was not statistically significant. Likewise, mean SW was significantly higher when anticipating a medio-lateral perturbation compared to when no perturbation was expected. People who are afraid of falling often show increased mean SW compared to those without fear of falling [25]; the increase in mean SW may also reflect increased anxiety. However, the possibility that anxiety is responsible for our findings is not supported by the observation that long-term SW variability significantly increased from the first to last perturbed-walking trials. Typically, anxiety decreases with exposure to balance perturbations [7], so we would have expected anxiety to *decrease* from the first to last perturbed-walking trials. In our study, state anxiety was only measured at the end of each trial block, so it is unclear whether anxiety levels changed from trial to trial. To better understand the relationship between spatio-temporal gait variability and anxiety, future research could collect anxiety measures during and/or following each trial.

There are conflicting findings in the literature as to whether treadmill walking is an accurate representation of overground walking. The constant treadmill speed and width may have limited the range of step lengths, widths, and times. Indeed, one study found that long- and short-term spatiotemporal variabilities were significantly lower on a treadmill compared to overground walking in young, healthy adults, suggesting that a treadmill induces invariant gait patterns [26]. In contrast, other studies found that, treadmill walking was quantitatively similar to overground walking [27,28]. Furthermore, participants’ performance was constrained by the characteristics of the treadmill. The treadmill kept moving after the perturbation, so participants did not have the option to stop walking to regain balance. The treadmill also eliminated optic flow that participants typically receive when walking overground. Lateral motion, which integrates visual input, is more sensitive to noise that may contribute to variability [29]. As such, the lack of sensory input may have impacted our observed levels of variability. However, a treadmill was beneficial for this study as it allowed us to collect enough steps to reliably estimate spatio-temporal variability and to impose perturbations using the motion platform.

ML COM velocity was estimated from a single marker, which has been found to be robust for walking [30]. However, participants’ responses to the perturbation were often accompanied by arm movements, which are not accounted for when estimating COM position and velocity using a single marker. The lack of additional markers, especially on the arms, poses a limitation in calculating the steps to ML COM velocity restabilization.

In conclusion, our findings suggest that young, healthy adults increase their long-term step width variability in anticipation of a perturbation. While this increased variability may allow for exploration of the environment, and reflect a healthy optimal range that allows for flexible and adaptable movement, it appeared to have no effect on ML COM velocity restabilization in the present study.

## Acknowledgements

We would like to thank Adam Lu and Jordan Oddo Silva for their assistance with data collection and processing.

## Notes

**Funding:** This work was supported by the Natural Sciences and Engineering Research Council of Canada (RGPIN-2014-04199) and the Ministry of Research and Innovation (Ontario). The authors acknowledge the support of the Toronto Rehabilitation Institute; equipment and space have been funded with grants from the Canada Foundation for Innovation, Ontario Innovation Trust, and the Ministry of Research and Innovation. Avril Mansfield was supported by a New Investigator Award from the Canadian Institutes of Health Research (MSH-141983). These funding sources had no role in the design or execution of this study, analyses or interpretation of the data, or decision to submit results.

### Competing Interest Statement

The authors have declared no competing interest.

